# Infection by a eukaryotic gut parasite in wild *Daphnia* sp. associates with a distinct bacterial community

**DOI:** 10.1101/2022.02.25.481906

**Authors:** Amruta Rajarajan, Justyna Wolinska, Jean-Claude Walser, Minea Mäder, Piet Spaak

**Author notes:** **Corresponding author email:** /.

## Abstract

Host-associated bacterial communities can play an important role in host fitness and resistance to diseases. Yet, few studies have investigated tripartite interaction between a host, parasite and host-associated bacterial communities in natural settings. Here, we use 16S amplicon sequencing to compare gut- and body-bacterial communities of wild water fleas belonging to the *Daphnia longispina* complex, between uninfected hosts and those infected with the common and virulent eukaryotic gut parasite *Caullerya mesnili* (Family: Ichthyosporea). We report community-level changes in host-associated bacteria with the presence of the parasite infection; namely decreased alpha diversity and increased beta diversity at the site of infection, i.e. host gut (but not host body). We also report decreased abundance of bacterial taxa proposed elsewhere to be beneficial for the host, and an appearance of taxa specifically associated with infected hosts. Our study highlights the host-microbiota-infection link in a natural system and raises questions about the role of host-associated microbiota in natural disease epidemics as well as the functional roles of bacteria specifically associated with infected hosts.

## Introduction

Host-associated bacterial communities play an important role in host health and the outcome of infectious disease. Several studies document host-associated bacteria’s role in increased colonization resistance against parasites (Libertucci & Young, 2019; Sekirov, Russell, Antunes, & Finlay, 2010). Host-associated microbiota may be beneficial to the host by contributing to the activation of the host’s adaptive immune response to the pathogen and/or competitively exclude pathogens by monopolizing host resources (Buffie & Pamer, 2013; Sorbara & Pamer, 2019).

However, bacterial community members that are beneficial or commensal to the host may also indirectly contribute to increased susceptibility to infectious disease. For instance, commensal members of the host-associated bacterial community that are generally present in the host in the absence of a pathogenic infection could expand into newly developed niches within the host upon disruption by a pathogen, hence switching to a pathogenic lifestyle (Bliska, Stevens, Bates, & King, 2021). The bacterial pathogen *Bacillus thuringiensis* is lethal to its gypsy moth host in the presence of a host-associated bacterial community, but not in antibiotic-treated hosts (Broderick, Raffa, & Handelsman, 2006) and similarly, germ-free hosts infected with the fungal pathogen *Beauveria bassiana* survive longer than infected hosts with microbiota (Wei et al., 2017). Thus, host-associated microbiota may either increase or decrease host susceptibility to infection by a pathogen. In general, a diseased state in the host may result from complex interactions between the host, associated microbes and the pathogen (Bass, Stentiford, Wang, Koskella, & Tyler, 2019; Bernardo-Cravo, Schmeller, Chatzinotas, Vredenburg, & Loyau, 2020; Pitlik & Koren, 2017).

There is now an increased appreciation for the putative role of bacterial communities in natural epidemics of infectious diseases (Stencel, 2021). Infection by a parasite leads to disease-induced dysbiosis or progressive changes in the host’s bacterial community structure, with late-stage infections typically correlating with a lower diversity of associated microbes (Chen, Ng, Wu, Chen, & Wang, 2017; Jani & Briggs, 2014; Lloyd & Pespeni, 2018; Nunez-Pons, Work, Angulo-Preckler, Moles, & Avila, 2018; Rosado et al., 2019) and an increased occurrence of opportunistic pathogens (Cornejo-Granados et al., 2017; Griffiths et al., 2019). An increase in dysbiosis-related bacteria in a host caused by a pathogenic infection (or an ecological stressor) may even increase susceptibility of a host to disease (Boutin, Bernatchez, Audet, & Derome, 2013; Gustin, Cromarty, Schifanella, & Klatt, 2021; Hinderfeld & Simoes-Barbosa, 2020). Hence, an understanding of host-associated microbial community changes in response to infectious disease epidemics has implications in wildlife disease management in both terrestrial (Allender, Baker, Britton, & Kent, 2018; Denman et al., 2018; Woodhams et al., 2014) and aquatic (Luter et al., 2017; Meyer et al., 2019; Quintanilla et al., 2018; Vezzulli, Pezzati, Huete-Stauffer, Pruzzo, & Cerrano, 2013)} systems.

Conversely, commensal host-associated bacterial communities may play a role in susceptibility to infections and consequent epidemics - though studies investigating this have been limited due to the challenging nature of inferring causal relations in the field. Soil microbiota dynamics influence the development of protistan clubroot disease in the Rutabega plant, *Brassica napus* (Daval et al., 2020). Populations of the common frog *Rana temporaria* that experienced repeated epidemics of *Ranavirus* had distinct bacterial communities compared to populations that did not (Campbell, Garner, Hopkins, Griffiths, & Harrison, 2019). Similarly, populations of the common Midwife toad that experience large, recurrent epidemics of the fungal pathogen *Batrachochytrium dendrobatidis* are associated with a distinct skin microbiota, which is not explained by distinct pathogen genotypes across sites (Bates et al., 2018). Further, epidemiological dynamics of *B. dendrobatidis* are linked with host-associated bacterial communities (Jani, Knapp, & Briggs, 2017; Longo, Savage, Hewson, & Zamudio, 2015).

The zooplankter water flea *Daphnia* is a keystone species in freshwater ecosystems (Lampert). It reproduces by cyclical parthenogenesis, with clonal reproduction being the dominant reproductive mode, and sexual reproduction occurring only under unfavourable environmental conditions (typically in late autumn). Two studies have investigated the role of *Daphnia-*associated bacterial communities during a parasite infection. Gut bacteria were found to play no role in *Daphnia magna* susceptibility to its bacterial pathogen *Pasteuria ramosa* (Sison-Mangus, Metzger, & Ebert, 2018). Also, germ-free *D. magna* that received microbiomes of *Daphnia* pre-exposed to a mixture of parasites did not show increased survival or tolerance upon re-exposure to the same parasites (Bulteel, Houwenhuyse, Declerck, & Decaestecker, 2021). Both studies investigated the influence of host-associated bacterial communities on *Daphnia* infection patterns under laboratory settings. However, no studies so far have investigated links between host-associated bacterial communities and parasite infections in wild *Daphnia* populations.

Populations of the *Daphnia longispina* complex in the lake Greifensee are infected with the highly virulent, eukaryotic gut parasite, *Caullerya mesnili* (Gonzalez-Tortuero et al., 2016; Wolinska, Bittner, Ebert, & Spaak, 2006; Wolinska, Keller, Bittner, Lass, & Spaak, 2004) (Family: Ichthyosporea) (Lu et al., 2020). Epidemics of *C. mesnili* (hereafter referred to as *Caullerya*) are seasonal and peak during late autumn and winter (Turko et al., 2018). Parasitization by *Caullerya* exerts a strong selection pressure on the host (Schoebel, Wolinska, & Spaak, 2010; Turko et al., 2018) by drastically reducing host fecundity and increasing mortality (Lohr, Laforsch, Koerner, & Wolinska, 2010). In this study, we investigate the relationship between infection by *Caullerya* and host-associated bacterial community composition in the *D. longispina* complex during a natural epidemic of the parasite. Specifically, we identify bacterial taxa that may uniquely associate with *Caullerya-*infected *Daphnia*. For this, we profiled bacterial communities of infected and uninfected hosts (in both their guts and body tissue) using 16S amplicon sequencing. We hypothesized that: (a) alpha diversity of gut bacterial communities would be lower in infected compared to uninfected hosts, consistent with pathogen-mediated dysbiosis (b) beta diversity of gut bacterial communities should differ between infected and uninfected hosts, reflecting colonization by putatively opportunistic bacterial pathogens and/or dominance of bacterial taxa related to dysbiosis in infected hosts and (c) differences in alpha and beta diversity of bacterial communities between infected and uninfected hosts should be limited to gut tissue (i.e. host body tissue should not differ in their bacterial communities by infection status) since the infection is specific to the gut.

## Material & Methods

### *Daphnia* sampling and processing

All *Daphnia* in the study were collected on 22/12/2020 from lake Greifensee (N 47°20′41″, E 8°40′21″) from a single sampling location using four vertical tows (0-30 m) with a 150μm mesh plankton net (Turko et al., 2018). The zooplankton sample was transferred to a 10L canister half-filled with surface lake water from the same location. In this study, we define a *Caullerya* ‘infection’ as visible, late-stage infection. A late stage *Caullerya* infection presents as spore clusters in the *Daphnia* gut, typically 8-12 days after initial exposure to the parasite (Lohr et al., 2010). For identifying *Caullerya-*infected *Daphnia*, zooplankton in the sample were collected on a ∼500μm mesh and *Daphnia* were visually screened for the presence of *Caullerya* spores in the gut under 40-50x magnification (Lohr et al., 2010). Ten adult *Daphnia* (>1mm in size) containing *Caullerya* spores in the gut were gently lifted by the antennae using sterilized forceps and placed by either PS or MM in a bottle containing 10mL Greifensee lake water filtered through a 0.45μm mesh. For uninfected *Daphnia*, 10 adult *Daphnia* that did not have visible *Caullerya* spore clusters in the gut were collected in the same way. No criteria other than the size cut-off, exclusion of individuals carrying resting eggs (indicating sexual reproduction) and presence/absence of *Caullerya* spores were used to exclude *Daphnia* from the study. Forceps were wiped with 10% bleach between every animal picked up to minimise cross-contamination of spores and/or bacteria. Sixteen bottles each of 10 infected and 16 bottles each of 10 uninfected *Daphnia* were prepared in alternating order. This was done to ensure that infected and uninfected *Daphnia* were processed from collection to dissection at the same average duration.

### Dissection and sample preparation

Sets of *Daphnia* picked into 10mL bottles (see above) were used for the preparations of gut bacterial and body bacterial community samples. For gut samples, 20 *Daphnia* per replicate were dissected under a stereo microscope, each in individual droplets of nuclease-free water using sterilized forceps (these 20 *Daphnia* originated from two bottles of 10 *Daphnia* prepared by PS and MM, to minimize systematic human error in identification of infected animals), resulting in eight replicates per treatment. Extracted guts were immediately pooled into a separate 20μL droplet of nuclease-free water, and afterwards transferred with a pipette to a 1.5mL microcentrifuge tube containing 10μL nuclease-free water. For paired body samples, the remaining tissue after extraction of guts was similarly pooled into a separate 1.5mL microcentrifuge tube. Forceps were wiped with 10% bleach between every dissection to minimize cross-contamination between *Daphnia* gut and body samples. All *Daphnia* were dissected within 24 hours of being collected from the lake. All samples were stored at -20°C immediately upon dissection until further processing. Four dissection negative controls (each containing 30μL of the water used to dissect *Daphnia*) were also prepared.

### Bacterial community profiling

DNA was extracted using the Qiagen Blood & Tissue kit (Cat #69506). Two DNA extraction blanks (tubes without biological material) were added to the workflow. All samples were lysed at 56°C for four hours and the extraction protocol provided by the manufacturer was followed. Samples were eluted in 40μL kit elution buffer for 20 min. A nested PCR approach was used to account for low biomass of our samples; at this stage, two PCR no-template controls were added to the workflow. Barcoded universal 16S primers 515F (5’-GTGCCAGCMGCCGCGGTAA-3’) (Caporaso et al., 2011) and m806R (5’-GGACTACNVGGGTWTCTAAT-3’) (Apprill, McNally, Parsons, & Weber, 2015) were used to amplify the V4-V5 region of the 16S gene using the following cycling conditions: initial denaturation at 95°C – 3 min; (98°C – 20 s, 52°C – 15 s, 72°C – 15 s) 32 cycles; followed by a final extension at 72°C for 5 min. This PCR step was done in triplicate for each sample, and triplicates were pooled before PCR purification. The location of all samples (including a total of eight negative controls) in the 96-well plates was randomized, though triplicates were always adjacent. PCR products were then purified using magnetic Agencourt AMPure XP beads (Cat #A63881) before being used in a second PCR with indexing primers. Indexes from the Illumina Nextera XT V2 Library Prep Kit were added to the samples via a PCR, under the conditions: initial denaturation at 95°C – 3 min; (95°C – 30 s, 55°C – 30 s, 72°C – 30 s) 9 cycles; followed by a final extension at 72°C for 5 min. Indexed PCR products were purified again using AMPure beads. The samples were then quantified using a Qubit HS Assay, normalized and pooled into a 1.5nM library for amplicon sequencing. Paired-end sequencing was carried out using MiSeq - 600PE v3 run kit with 10% PhiX.

### Pre-processing of reads

Sequencing resulted in a total of 11.6M paired-end reads (minimum = 204 312, maximum = 519 591 per biological sample). Initial pre-processing steps were performed on the Euler computing cluster at ETH Zürich. Raw reads were 5’-end trimmed, merged, and quality filtered (see supplementary file for more details). Amplicons were clustered using UPARSE2 (Edgar, 2013), denoised into zero-radius Operational Taxonomic Units (ZOTUs) using UNOISE3 (Robert C. Edgar, 2016), further clustered based on 97% sequence similarity and annotated using non-Bayesian SINTAX classifier v.138 (R. C. Edgar, 2016) and the Silva database (Quast et al., 2013). All subsequent steps were performed in R v4.0.2 using the package phyloseq (McMurdie & Holmes, 2013). 29 ZOTUs (14 of unidentified phyla, 13 chloroplasts, 2 mitochondria) were filtered from the dataset, resulting in a total of 737 ZOTUs. ZOTUs discovered in negative controls were not removed, as this was not suitable for our dataset; see Fig S1 and Table S1 for a detailed description of negative controls and their analyses and also (Hornung, Zwittink, & Kuijper, 2019; Kim et al., 2017). After exclusion of samples that failed sequencing (<200 reads), the following number of samples remained in the study: 8 samples of infected guts, 7 of infected bodies, 7 of uninfected guts and 7 of uninfected bodies, representing a total of 160 *Caullerya-*infected and 140 uninfected individuals. All samples were then rarefied to an even depth of 170 000 reads. The SRS method of normalizing library sizes (Beule & Karlovsky, 2020) yielded the same statistical results as rarefaction and hence only the results of the rarefied dataset are presented (see below). The rarefaction step resulted in the removal of 54 ZOTUs, leaving 683 ZOTUs in the dataset.

### Statistical analyses

All statistical analyses were done in R v4.0.2 using the phyloseq package. First, the dominant bacterial orders were visualised. ZOTUs were aggregated at the order level using the *tax_glom* function. Rare orders were lumped into the category “Other” (i.e. orders constituting <1% of biological samples and not present in every sample). Differential abundance of dominant bacterial orders was tested between *Caullerya-*infected and uninfected *Daphnia*, separately for gut and body bacterial communities using the Wald test of the DESeq2 package (Love, Huber, & Anders, 2014) and corrected for multiple comparisons using the “fdr” method.

Second, diversity indices were compared between *Caullerya-*infected and uninfected *Daphnia*, separately for gut and body tissue since we were primarily interested in the effects of infection. All diversity analyses were performed at the ZOTU level. Alpha-diversity measures (qualitative: ZOTU richness and quantitative: Inverse Simpson Index) of samples were estimated using the *estimate_richness* function. One-way ANOVAs were performed on both alpha diversity measures to test for variation between infected and uninfected *Daphnia*. Next, beta diversity indices (qualitative: Jaccard dissimilarity and quantitative: weighted unifrac distance) were visualised using PCOA plots. One-way PERMANOVAs were performed using the *adonis* function of the vegan package (9999 perm). PERMANOVAs of beta diversity metrics were only performed after checking if the data meets the homogeneity of dispersion assumption using the *betadisper* function of the vegan package. Percentage variation explained by the models used for beta diversity analyses was estimated with a db-RDA using the *capscale* function.

Finally, to identify ZOTUs indicative of *Daphnia* infection status, the *signassoc* function of the Indicspecies package in R was used (two-tailed test, 9999 perm, corrected for multiple comparisons using the Sidak method) (De Caceres & Legendre, 2009). This was not done separately for gut and body tissue since we aimed to identify bacterial taxa that associate significantly with infected (or uninfected) *Daphnia* regardless of host tissue. The relative abundances of indicator ZOTUs were then visualised in a heatmap using ggplot2.

## Results

### Bacterial community composition

Gut and body bacterial communities of *Daphnia* sampled from lake Greifensee comprised of 18 dominant orders (Fig. 1). Bodies of infected *Daphnia* had a higher relative abundance of Enterobacteriales (18.21 ± 9.23%, mean ± SD) and Pseudomonadales (14.37 ± 24.47%) compared to bodies of uninfected *Daphnia* (3.54 ± 2.58% and 10.29 ± 22.58%, respectively). Micrococcales were rare within *Daphnia;* but were significantly more abundant in uninfected bodies (0.009 ± 0.006%) compared to infected bodies (0.005 ± 0.003%). The above bacterial orders that differed by infection status in *Daphnia* bodies followed a similar (although not significant) trend in *Daphnia* guts. Enterobacteriales constituted 38.9 ± 24.77% of *Caullerya-*infected guts compared to 11.7 ± 7.45% of uninfected guts, while Micrococcales formed 0.027 ± 0.034% of uninfected guts compared to 0.014 ± 0.009% of *Caullerya-*infected guts (Fig.1). Nevertheless, bacterial community composition varied substantially across replicates; two gut-body pairs (one *Caullerya-*infected and one uninfected) were dominated by the order Pseudomonadales, which was typically less abundant in all other samples (see Tables S2 and S3). Moreover, one *Caullerya-*infected gut sample (replicate 1 in top, left panel of Fig. 1) unexpectedly showed a bacterial community composition more similar to uninfected *Daphnia* guts. We repeated the analysis of differential abundance in dominant bacterial orders after removing this sample and found that the order Rickettsiales was significantly more abundant in uninfected (36.36 ± 12.89%) compared to infected (12.84 ± 6.66%) *Daphnia* guts (W = 3.83, *p*adj *=* 0.01, data not shown).

**Fig. 1.**
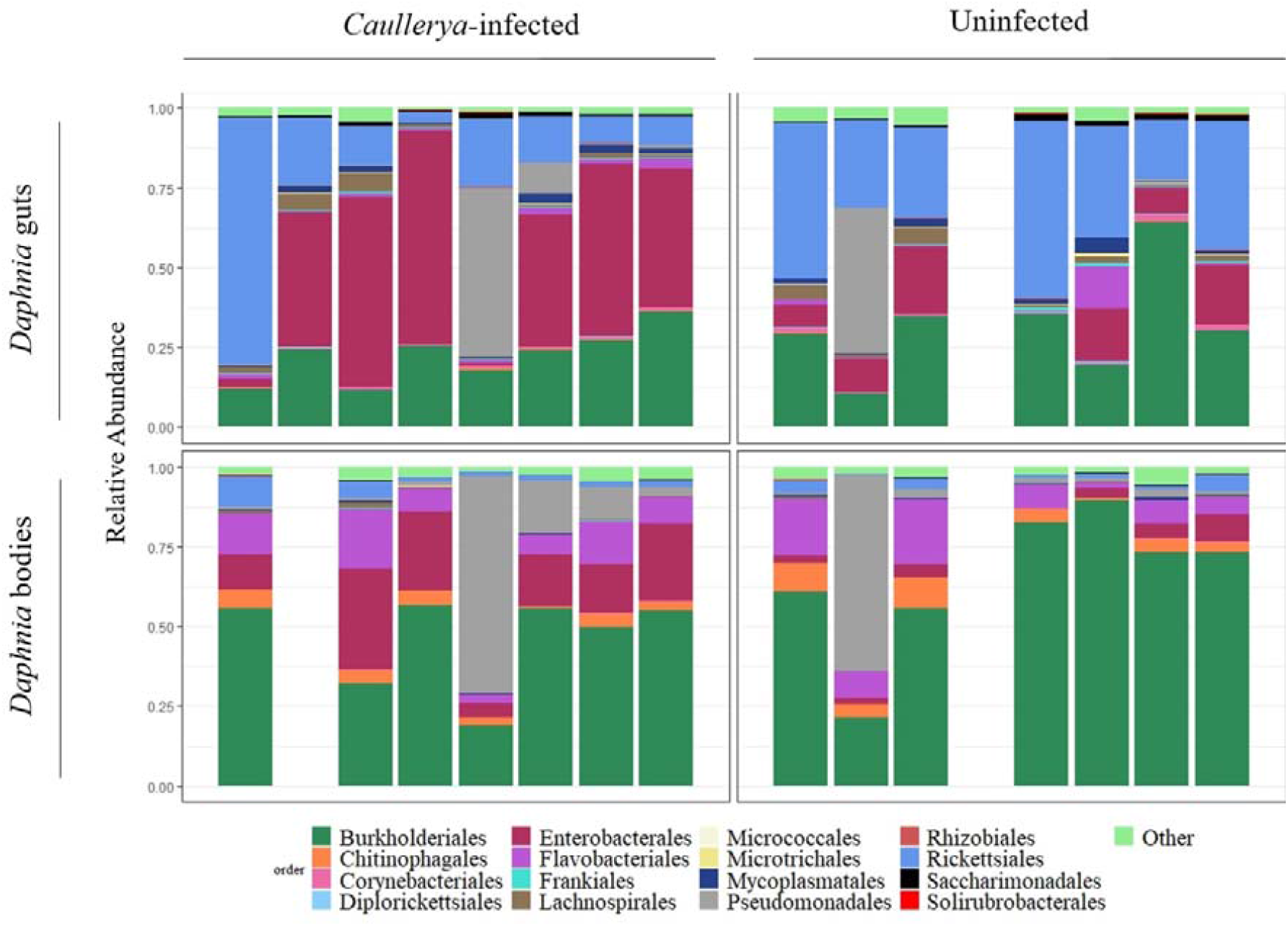
Bacterial communities in *Caullerya-*infected (left panels) and uninfected (right panels) *Daphnia. Daphnia* gut communities are shown on the top and body communities at the bottom. Rare taxa, i.e. bacterial orders comprising <1% of the total dataset and not present in every sample were classified as “Other”, containing 90 orders. Empty spaces correspond to samples that failed sequencing and hence were not part of analysis.

### Alpha diversity

We used ZOTU richness (qualitative) and Inverse Simpson Index (quantitative) measures of alpha diversity. ZOTU richness did not vary between *Caullerya-*infected and uninfected *Daphnia* regardless of host tissue (Fig. 2). The Inverse Simpson Index was significantly lower in *Caullerya-*infected guts compared to uninfected guts (F = 6.2, *p =* 0.027); however, *Daphnia* bodies showed an opposite (although not significant) trend (Table 1).

**Table 1.** One-way ANOVAs of alpha diversity metrics ZOTU richness (A) and Inverse Simpson Index (B) across *Caullerya*-infected and uninfected *Daphnia* separately for *Daphnia* gut and body samples. Model used was diversity metric ∼ infection. \**p* < 0.05

|  |  |  |  |  |  |  |
| --- | --- | --- | --- | --- | --- | --- |
| (A) **ZOTU Richness** |  |  |  |  |  |  |
|  |  | Df | Sum Sq | Mean Sq | F value | *P* value |
| **gut** | infection | 1 | 3320 | 3320 | 1.56 | 0.23 |
|  | Residuals | 13 | 27655 | 2127 |  |  |
|  |  | Df | Sum Sq | Mean Sq | F value | *P* value |
| **body** | infection | 1 | 2498 | 2498 | 0.83 | 0.38 |
|  | Residuals | 12 | 36270 | 3022 |  |  |
| (B) **Inverse Simpson Index** |  |  |  |  |  |  |
|  |  | Df | Sum Sq | Mean Sq | F value | *P* value |
| **gut** | infection | 1 | 10.5 | 10.46 | 6.2 | **0.027\*** |
|  | Residuals | 13 | 21.9 | 1.69 |  |  |
|  |  | Df | Sum Sq | Mean Sq | F value | *P* value |
| **body** | infection | 1 | 11.3 | 11.3 | 1.95 | 0.19 |
|  | Residuals | 12 | 69.6 | 5.8 |  |  |

**Fig. 2.**
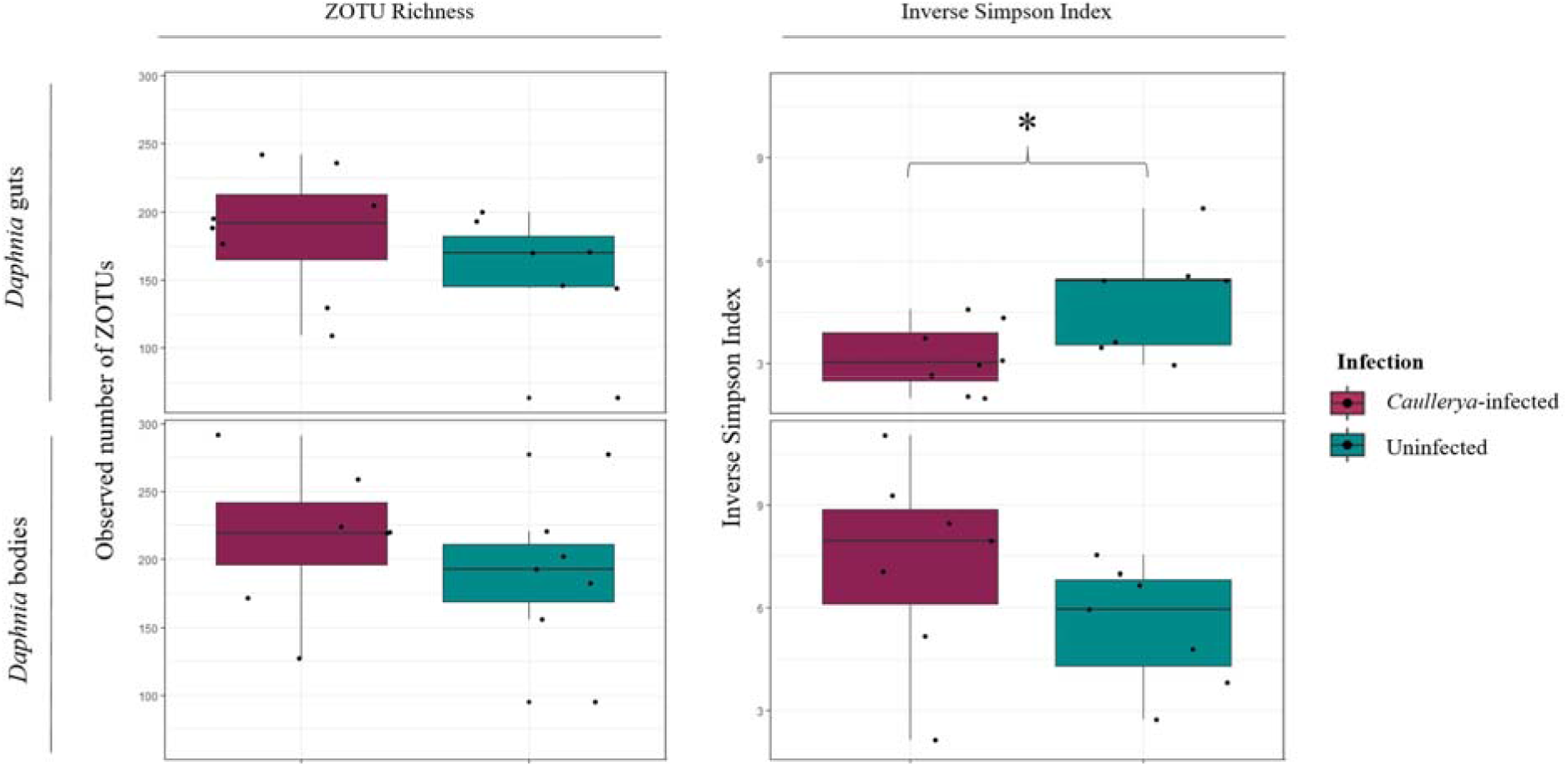
Alpha diversity indices: ZOTU richness (left panels) and Inverse Simpson Index (right panels) in *Caullerya-*infected (maroon) and uninfected (green) *Daphnia*-associated bacterial communities. Panels are indicated with tissue type; with *Daphnia* guts on top panels and *Daphnia* body in bottom panels, for each metric. \**p* < 0.05, one-way ANOVA.

### Beta diversity

We used the Jaccard Dissimilarity (qualitative) and Weighted Unifrac distance (quantitative) measures of beta diversity. Jaccard Dissimilarity which is a count of the number of unshared taxa, differed significantly between *Caullerya-*infected and uninfected *Daphnia*, in both the guts (Fig. 3a, F = 3.01, *p* = 0.0093) and bodies (Fig. 3b, F = 2.00, *p* = 0.031). The Weighted Unifrac distance which weighs differences in ZOTUs by their relative abundance and phylogenetic relatedness was significantly different between *Caullerya-*infected and uninfected *Daphnia* in the guts (Fig. 3b, F = 3.115, *p* = 0.0386), but not the *Daphnia* bodies (Fig. 3d, Table 2). This implies that *Caullerya-*infected *Daphnia* in general have phylogenetically distinct set of bacteria compared to uninfected *Daphnia*, but that they are differentially abundant only at the site of *Caullerya* infection – i.e. in the gut and not the body tissue.

**Table 2.** One-way PERMANOVAs of beta diversity metrics Jaccard Dissimilarity (A) and Weighted Unifrac Distance (B) across *Caullerya*-infected and uninfected *Daphnia* (9999 permutations). Analyses were conducted separately for each tissue type, indicated on the left. %Variation column shows the % variation explained by the model using db-RDA. \**p* < 0.05

| (A) |  | Jaccard Dissimilarity |  |  |  |  |  | %Variation |
| --- | --- | --- | --- | --- | --- | --- | --- | --- |
|  |  | Df | SumsOfSqs | MeanSqs | F.Model | R2 | Pr(>F) |  |
| gut | infection | 1 | 0.59 | 0.59 | 3.01 | 0.188 | **0.0093\*** | 18.8 |
|  | Residuals | 13 | 2.55 | 0.196 |  | 0.812 |  |  |
|  | Total | 14 | 3.14 |  |  | 1 |  |  |
|  |  | Df | SumsOfSqs | MeanSqs | F.Model | R2 | Pr(>F) |  |
| body | infection | 1 | 0.365 | 0.365 | 2 | 0.143 | **0.031\*** | 14.3 |
|  | Residuals | 12 | 2.192 | 0.183 |  | 0.857 |  |  |
|  | Total | 13 | 2.556 |  |  | 1 |  |  |
| (B) |  | Weighted Unifrac Distance |  |  |  |  |  | %Variation |
|  |  | Df | SumsOfSqs | MeanSqs | F.Model | R2 | Pr(>F) |  |
| gut | infection | 1 | 0.848 | 0.848 | 3.115 | 0.193 | **0.0386\*** | 19.4 |
|  | Residuals | 13 | 0.354 | 0.027 |  | 0.803 |  |  |
|  | Total | 14 | 0.437 |  |  | 1 |  |  |
|  |  | Df | SumsOfSqs | MeanSqs | F.Model | R2 | Pr(>F) |  |
| body | infection | 1 | 0.028 | 0.028 | 1.694 | 0.124 | 0.1536 | 12.5 |
|  | Residuals | 12 | 0.197 | 0.017 |  | 0.876 |  |  |
|  | Total | 13 | 0.225 |  |  | 1 |  |  |

**Fig. 3.**
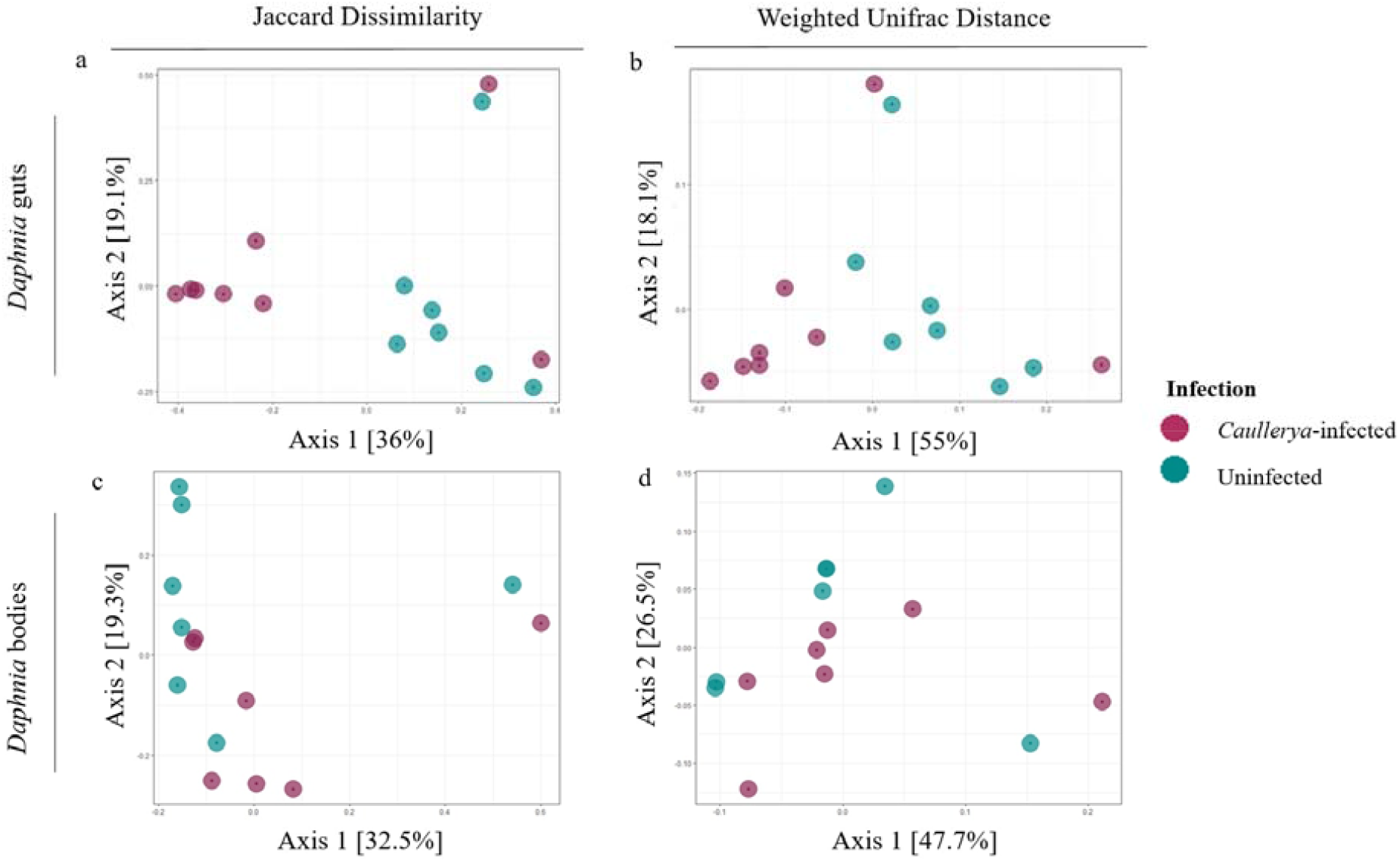
PCOA plots of beta diversity measures of *Daphnia* gut bacterial communities (top panels) and *Daphnia* body bacterial communities (bottom panels). Jaccard Dissimilarity (Fig. 3a and 3c) and Weighted Unifrac Distance (Fig. 3b and 3d); colors represent *Caullerya-*infected (maroon) and uninfected (green) bacterial communities.

### Indicator taxa

Indicator analysis pointed to 10 ZOTUs for which abundance distributions could identify *Caullerya* infection status of *Daphnia* samples, and included two ZOTUs that were highly abundant in the dataset. A majority of the discovered indicator ZOTUs (9 out of 10) were more abundant in *Caullerya-*infected than in uninfected *Daphnia* (Fig. 4, Table S4).

**Fig. 4.**
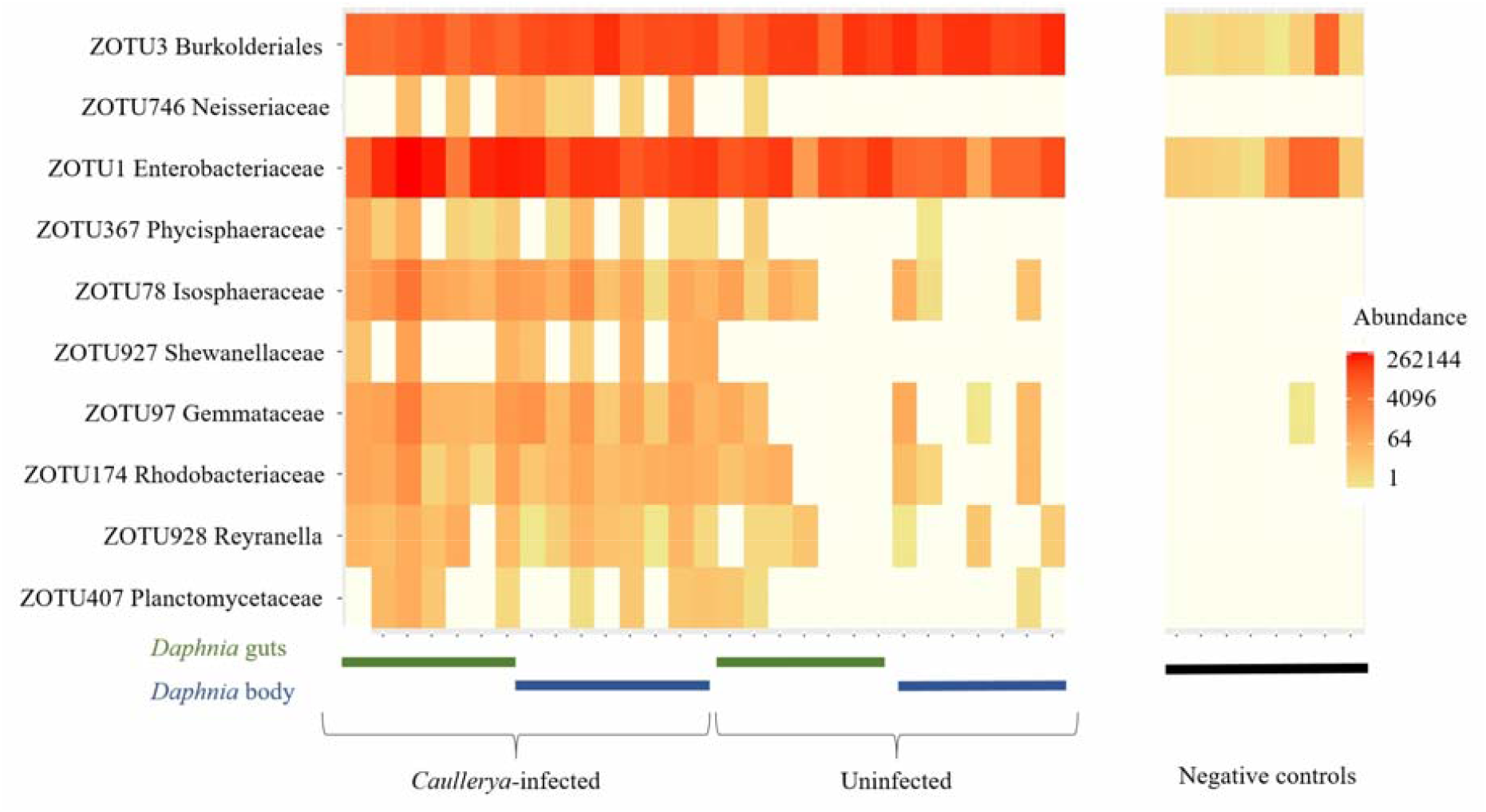
Heatmap of log-transformed ZOTU counts, showing abundance of ZOTUs that indicate presence or absence of *Caullerya* infection in *Daphnia* samples, as identified by Indicspecies. Rows correspond to ZOTUs (with only the family shown) and columns represent infected or uninfected samples (*Daphnia* gut, *Daphnia* body or negative control, as indicated. See methods for details on negative controls.)

## Discussion

In this study, we compared the bacterial communities of gut and body tissue between *Caullerya-*infected and uninfected *Daphnia* in the wild. We found that host gut bacterial communities were significantly less even (as indicated by a lower Shannon Index), confirming our hypothesis that alpha diversity of the gut bacterial community is lower in infected individuals and suggesting an abundance shift of some bacteria. Further, gut bacterial communities of infected and uninfected hosts also differed in their beta diversity indices, confirming our hypothesis of differing beta diversity by infection status and indicating more variable bacterial communities in the presence of an infection. However, we also hypothesized that the above-mentioned changes in bacterial community diversity would apply only to host gut but not body tissue, since in the investigated infection is gut-specific. For this, we obtained mixed results: gut bacterial communities varied in both their composition and relative abundance - but an infection in the gut also correlated with distinct community composition in the host body tissue. Hence, bacterial communities in the host body too are altered in the presence of a gut infection. Further, infected and uninfected hosts had distinct associated ZOTUs. Overall, these patterns are consistent with changes in host-associated bacterial communities concurrent with infection in other wild animals such as mallards (Ganz et al., 2017) and bats (Wasimuddin et al., 2018).

In the present study, the visual presence of parasite spores in the host gut is an indicator of late-stage infection; fully mature spores in the gut are visible 8-12 days post initial exposure to the parasite (Lohr et al., 2010). Therefore, during an ongoing epidemic in the field, the hosts classified as uninfected in our study could include individuals with undetected earlier stage infections. Nevertheless, we find that the gut bacterial communities of infected hosts with visible parasite spore clusters are distinct in alpha and beta diversity from those of uninfected and/or early stage infected hosts. This suggests that the infection most likely preceded the formation of distinct bacterial communities compared to uninfected hosts.

A previous experiment in *Daphnia galeata* reported a suppression of immune-related gene expression through a downregulation in C-type lectins, which play a role in pathogen cell recognition, 48 hours after exposure to *Caullerya* (Lu et al., 2018). C-type lectin expression is intricately linked with host microbiota; in the kuruma shrimp *Marsupenaeus japonicus*, C-type lectin expression regulates gut bacterial community composition (Zhang, Zhang, & Wang, 2021). Conversely, intestinal microbiota epigenetically downregulate C-type lectin expression in the mouse host to prevent adherence by a gut pathogen (Woo et al., 2019). We speculate that such a downregulation of immune-related genes in the host in the present study may also contribute to an increase in ZOTU richness associated with the bacterial communities of infected *Daphnia* bodies (Fig. 1), since more bacteria may colonize hosts with a suppressed immune response.

Our result does not eliminate the possibility of reverse directionality, i.e. associated bacteria also playing a role in susceptibility to infection. Epidemics of infectious diseases are often triggered by the host population experiencing an ecological stressor (Gehman, Hall, & Byers, 2018; LaDeau, Allan, Leisnham, & Levy, 2015; Lafferty & Holt, 2003). For example, salinity stress in *Rana sylvatica* compromises healthy amphibian immune response against *Ranavirus* and leads to greater transmission of the pathogen as well as higher disease-induced host mortality (Hall, Brunner, Hutzenbiler, & Crespi, 2020). Further, environmental stressors may indirectly impact population health by altering host-associated bacterial communities (Greenspan et al., 2020). Outbreaks of *Caullerya* in wild *Daphnia* populations correlate with cyanobacterial blooms, as confirmed with field observations and lab experiments (Tellenbach et al., 2016). Cyanobacterial blooms are associated with dramatic changes in the structure of lacustrine pelagic bacterial communities (Tromas et al., 2017). On the other hand, freshwater zooplankton such as *Daphnia* undergo considerable shifts in associated bacterial communities driven by changes in environmental bacterial communities in general (Callens, De Meester, Muylaert, Mukherjee, & Decaestecker, 2020; Eckert, Anicic, & Fontaneto, 2021) and by exposure to cyanobacteria in particular (Macke, Callens, De Meester, & Decaestecker, 2017). At the time of *Daphnia* collection for the present study, the lake had a large bloom of cyanobacteria, particularly of the genus *Gomphosphaeria* (unpublished data). We speculate that a cyanobacterial diet or an altered pelagic bacterial community concurrent with cyanobacterial blooms may have initiated a perturbation in host bacterial communities, predisposing them to infection and the infection then further altered bacterial community structure.

The bacterial community structure of infected and uninfected hosts at the order level also points to physiological stress in the host population in the present study. An overgrowth of the family Enterobacteriaceae is associated with intestinal inflammation and dysbiosis in mammals including mice and humans (Chong, Cheng, Hogg, & Belov, 2020; Lupp et al., 2007; Zeng, Inohara, & Nunez, 2017). Though the family Enterobacteriaceae (particularly ZOTU1, Family: Enterobacteriaceae) was more abundant in infected hosts (29.00 ± 21.38%), it was also among the dominant taxa in uninfected hosts (8.00 ± 6.8%). Enterobacteriaceae was described as a core taxon in rotifers and crustaceans including *Daphnia* in one study (Eckert et al., 2021), but other studies have showed it was typically rare (Akbar et al., 2021; Callens, Watanabe, Kato, Miura, & Decaestecker, 2018; Macke et al., 2020) or absent (Freese & Schink, 2011) in laboratory-cultured, healthy *Daphnia* guts. Thus, the increased abundance of this order suggests a stressed host population independent of infection, possibly mediated by blooms of cyanobacteria in the lake.

Changes in bacterial community structure, particularly an increased prevalence of opportunistic pathogens, have been suggested to interact with the primary causative agent of a disease to produce a diseased state in the host (Bernardo-Cravo et al., 2020; Egan & Gardiner, 2016). In the present study, infected hosts contained differentially abundant ZOTUs compared to uninfected *Daphnia*. Several of these are relatively unknown aquatic microbes; hence apart from their occasional association with aquatic hosts, very little is known about their functional ecology or pathogenic potential (see Table S5). The ZOTUs significantly more abundant in infected hosts contain metabolically diverse species; several of the highlighted bacterial genera or families e.g. ZOTU927 (Genus: *Shewanella*), ZOTU174 (Family: Rhodobacteraceae) and ZOTU97 and ZOTU78 (Phylum: Planctomycetes) contain both opportunistic pathogens and symbionts with probiotic properties aiding disease resistance in a range of hosts. ZOTU3 (Family: Burkholderiales), which showed an increased abundance in uninfected *Daphnia* is a documented beneficial symbiont of *Daphnia* (Cooper & Cressler, 2020; Peerakietkhajorn, Kato, Kasalicky, Matsuura, & Watanabe, 2016) and is consistently reported to be among the most abundant host-associated bacterium across *Daphnia* species (Qi, Nong, Preston, Ben-Ami, & Ebert, 2009).

One aspect of host-parasite-bacterial community interaction that remains unaddressed in our study is the role of the host genotype in these interactions. A comprehensive study in sticklebacks showed that exposure to a parasitic helminth initially caused divergent, host genotype-specific changes in gut bacterial communities; however, successfully infected hosts converged in bacterial community composition across host genotypes (Hahn et al., 2022). The *Daphnia – Caullerya* host-parasite system is characterized by genetic specificity of infection (Turko et al., 2018; Yin, Petrusek, Seda, & Wolinska, 2012). However, the extent of host genotype influence in the assembly of *Daphnia-*associated bacterial communities is still debated, with conflicting results in laboratory studies (Callens et al., 2020; Frankel-Bricker, Song, Benner, & Schaack, 2020; Sullam, Pichon, Schaer, & Ebert, 2018). As such, zooplankton including *Daphnia* are proposed to be open systems with their bacterial communities showing high flexibility to changes in environmental bacterial composition (Eckert et al., 2021; Mushegian, Walser, Sullam, & Ebert, 2018). Further, variation in *Daphnia-*associated bacterial communities by genotype in natural settings is unknown. Since we did not genotype the hosts in the present study, it is possible that the pools of infected and uninfected samples comprised of partially different genotypes – but the extent of host genotype influence in these interactions requires further investigation. Thus, further studies correlating host genotype, infection by a parasite and bacterial community structure in the wild would be required to fully disentangle the complex effects of various factors that influence host bacterial community structure. Nevertheless, studies like the present one, which investigate the bacterial communities of wild hosts experiencing multiple environmental stressors, could provide insight into the ecological and evolutionary significance of host-associated microbiota.

## Acknowledgement

We would like to thank the Spaak group, the department of Aquatic Ecology at the Swiss Federal Institute of Aquatic Science and Technology (Eawag) Dübendorf for helpful discussions during experiment design and data analysis and the Genetic Diversity Centre, ETH Zürich for additionally providing extensive support during library preparation and sequencing. We also thank Christoph Walcher for help collecting zooplankton samples from the lake. This work was supported by a joint “lead agency” grant from the German Science Foundation (WO 1587/6-1 to JW) and Swiss National Science Foundation (310030 L 166628 to PS).

## Data Accessibility & Benefit Sharing Statement

### Data Accessibility Statement

Raw 16S sequence data have been submitted to DDBJ (https://www.ddbj.nig.ac.jp/index-e.html); an Accession ID will provided as soon possible. All processed data (OTU table, associated files) and R code to reproduce figures and tables in the manuscript will be made available on https://doi.org/10.25678/0005Z6 upon publication.

### Benefit Sharing Statement

Not applicable.

## Author Contributions

AR, JW and PS conceived and designed the study. AR, MM and PS performed experimental work. AR and JCW performed data analysis with inputs from JW and PS. AR wrote the original draft of the manuscript; JW edited the manuscript with inputs from all co-authors.

