## Supplementary material for "Infection by a eukaryotic gut parasite in wild *Daphnia* sp. associates with a distinct bacterial community"

**Analysis of negative controls:** “blank” samples were added at several stages of the workflow (see Methods): i.e. we had four dissection blanks, two DNA extraction blanks and two PCR no-template controls, resulting in eight negative controls against 32 experimental samples. Three of the eight negative controls had <100 reads (two dissection negatives and one PCR negative), while the remaining five had between 16 853 and 240 206 reads (compared to biological samples which had between 170 000 and 518 570 reads). Due to this large variation in the number of reads among negative controls, we deduced that contamination in this dataset arose from cross-contamination between samples as opposed to systematic contamination, since systematic contamination would result in all negative controls having a substantial number of reads. Further, we conclude that the cross-contamination likely occurred from biological samples to negative controls due to the following observations:

(a) The biological samples had a total of 683 ZOTUs, whereas the negative controls had 180 ZOTUs, which were a subset of the ZOTUs in the biological samples.

(b) The most dominant bacterial orders in biological samples (Fig. 1, e.g. Burkholderiales, Enterobacteriales, and Pseudomondales) were more abundant in biological samples than in negative controls (Table S1). These orders were also present in all negative controls. On the other hand, Order composition of negative controls showed that they were dominated by orders that were rare in biological samples – e.g. “Other”, i.e. orders that constituted <1% of, and were not present in all biological samples (Fig. S1). The orders Corynebacterales and Rhizobiales (also present but not dominant among biological samples) were similarly more abundant in negative controls than the biological samples, but these were absent in some negative controls (2/8 and 3/8 respectively). From this, we conclude that experimental samples had a biological signal, whereas negative controls comprised mostly of overamplified rare ZOTUs that likely originated from biological samples.

| **Order** | **% Relative abundance among biological samples (mean ± SD)** | **% Relative abundance among negative controls (mean ± SD)** |
| --- | --- | --- |
| **Burkholderiales** | 41.78 ± 22.50 | 10.25 ± 4.79 |
| **Corynebacteriales** | 0.47 ± 0.52 | 6.53 ± 6.12 |
| **Enterobacteriales** | 18.8 ± 19.27 | 5.88 ± 4.91 |
| **Pseudomonadales** | 9.81 ± 19.84 | 16.91 ± 5.12 |
| **Rhizobiales** | 0.06 ± 0.04 | 4.66 ± 5.30 |
| **Other** | 2.89 ± 1.3 | 55.77 ± 9.27 |

**Table S1** Percentage relative abundance of orders detected in negative controls versus biological samples. Orders that were absent in negative controls are not included.

(c) The most abundant ZOTUs among experimental samples were detected in all negative controls whereas rarer ZOTUs in experimental samples only occurred in a subset of negative controls. Abundant ZOTUs are the most prone to being cross-contaminated between samples; further, low biomass samples (such as negative controls) that are processed simultaneously with experimental samples are the most sensitive to receiving cross-contamination (1, 2). We verified that infected and uninfected samples in our study did not differ significantly in their biomass (i.e. DNA concentration immediately after DNA extraction, data not presented).

Since we suspected that ZOTUs in the negative controls likely originated from experimental samples, we chose not to filter them out of the dataset. Further, since abundance patterns across experimental and negative control samples varied (i.e. abundant ZOTUs in experimental samples were less abundant in negative controls and rare ZOTUs in experimental samples showed the opposite trend), we could not apply a rule to ‘subtract’ bacterial counts which would be consistent for all ZOTUs (3, 4).

Of the 10 ZOTUs identified as indicator ZOTUs in this study (main text Fig. 4), seven were absent in the negative controls. The other three showed the following prevalence pattern across negative control samples: ZOTU1 (8/8), ZOTU3 (8/8) and ZOTU97 (1/8), all of which were more abundant in biological samples than in negative controls.

The location of samples on the 96-well plate during library preparation was randomized (see Methods). Therefore, cross-contamination in our dataset was equally likely to occur within and between the groups formally compared in this study. Together with the above-mentioned patterns, we argue that the presence of ZOTUs in negative controls does not invalidate the detected statistical differences between infected and uninfected groups.

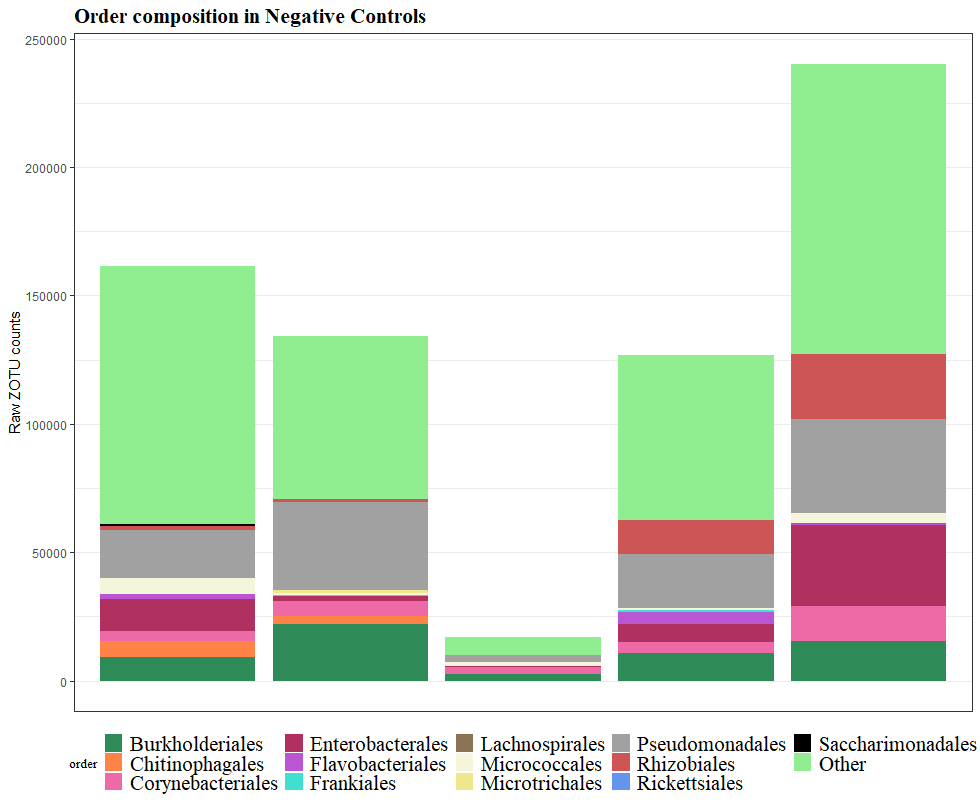

**Figure S1** Order composition (raw ZOTU count data) of negative control samples. Negative controls with <100 reads are not shown. “Other” corresponds to the same orders that were defined as “Other” in experimental samples (See Fig. 1), i.e. bacterial orders that formed <1% of the dataset and did not occur in every sample. Orders Diplorickettsiales, Mycoplasmatales and Solirubrobacterales present in Fig. 1 were absent in all negative controls.

|  | ***Daphnia* guts (uninfected vs. *Caullerya*)** | | |  | ***Daphnia* body (uninfected vs. *Caullerya*)** | | |
| --- | --- | --- | --- | --- | --- | --- | --- |
|  | log2FoldChange | Wald stat | padj |  | log2FoldChange | Wald stat | padj |
| **Burkholderiales** | 0.40 | 0.86 | 0.55 |  | 0.89 | 2.13 | 0.13 |
| **Chitinophagales** | -1.33 | -1.47 | 0.30 |  | 0.87 | 2.08 | 0.13 |
| **Corynebacteriales** | 0.42 | 0.81 | 0.55 |  | 0.24 | 0.59 | 0.67 |
| **Diplorickettsiales** | 1.60 | 2.81 | 0.08 |  | -0.12 | -0.27 | 0.78 |
| **Enterobacteriales** | -1.87 | -2.41 | 0.09 |  | **-1.96** | **-3.06** | **0.01*** |
| **Flavobacteriales** | -1.01 | -2.09 | 0.13 |  | 0.45 | 0.90 | 0.53 |
| **Frankiales** | 0.51 | 1.18 | 0.45 |  | -0.41 | -1.06 | 0.53 |
| **Lachnospirales** | -0.02 | -0.04 | 0.10 |  | -0.64 | -0.98 | 0.53 |
| **Micrococcales** | 1.08 | 1.57 | 0.30 |  | **1.55** | **3.16** | **0.01*** |
| **Microtrichales** | 0.19 | 0.43 | 0.81 |  | -0.45 | -1.31 | 0.40 |
| **Mycoplasmatales** | -0.01 | -0.06 | 0.98 |  | 0.83 | 1.58 | 0.32 |
| **Other** | 0.72 | 1.50 | 0.30 |  | 0.37 | 1.32 | 0.40 |
| **Pseudomonadales** | -2.29 | -2.52 | 0.09 |  | **-3.14** | **-3.42** | **0.01*** |
| **Rhizobiales** | -1.22 | -2.06 | 0.13 |  | -0.33 | -0.89 | 0.53 |
| **Rickettsiales** | 0.61 | 1.05 | 0.50 |  | -0.17 | -0.32 | 0.78 |
| **Saccharimonadales** | 0.43 | 0.87 | 0.55 |  | -0.24 | -0.53 | 0.67 |
| **Solirubrobacterales** | 0.16 | 0.28 | 0.89 |  | 0.21 | 0.71 | 0.63 |

**Table S2** Wald test for differential abundance of bacterial orders in infected vs uninfected *Daphnia* (done separately for gut and body samples). *p-*values were adjusted for multiple comparisons using the “fdr” method. Significant **p-*adj <0.05 are shown in bold.

|  | ***Daphnia* gut** | |  | ***Daphnia* body** | |
| --- | --- | --- | --- | --- | --- |
| **Order** | ***Caullerya-*infected** | **Uninfected** |  | ***Caullerya-*infected** | **Uninfected** |
| **Burkholderiales** | 22.208 ± 8.261 | 31.973 ± 16.695 |  | 46.125 ± 14.749 | 65.159 ± 22.541 |
| **Chitinophagales** | 0.303 ± 0.248 | 0.114 ± 0.132 |  | 3.592 ± 1.642 | 5.016 ± 3.076 |
| **Corynebacteriales** | 0.658 ± 0.215 | 0.951 ± 0.769 |  | 0.138 ± 0.095 | 0.12 ± 0.088 |
| **Diplorickettsiales** | 0.068 ± 0.033 | 0.227 ± 0.271 |  | 0.023 ± 0.017 | 0.017 ± 0.012 |
| **Enterobacteriales** | 38.898 ± 24.769 | 11.695 ± 7.449 |  | **18.209 ± 9.231** | **3.538 ± 2.581** |
| **Flavobacteriales** | 1.393 ± 0.925 | 2.406 ± 4.732 |  | 9.959 ± 5.465 | 9.669 ± 6.789 |
| **Frankiales** | 0.298 ± 0.146 | 0.485 ± 0.368 |  | 0.086 ± 0.075 | 0.045 ± 0.03 |
| **Lachnospirales** | 2.159 ± 1.989 | 2.303 ± 1.779 |  | 0.516 ± 0.614 | 0.192 ± 0.201 |
| **Micrococcales** | 0.014 ± 0.009 | 0.027 ± 0.034 |  | **0.005 ± 0.003** | **0.009 ± 0.006** |
| **Microtrichales** | 0.235 ± 0.090 | 0.321 ± 0.273 |  | 0.077 ± 0.089 | 0.035 ± 0.017 |
| **Mycoplasmatales** | 1.614 ± 1.188 | 1.722 ± 1.625 |  | 0.175 ± 0.101 | 0.267 ± 0.3 |
| **Other** | 2.073 ± 1.104 | 3.189 ± 1.465 |  | 3.267 ± 1.148 | 3.174 ± 1.385 |
| **Pseudomonadales** | 8.087 ± 18.409 | 6.762 ± 17.145 |  | **14.373 ± 24.472** | **10.291 ± 22.575** |
| **Rhizobiales** | 0.119 ± 0.041 | 0.059 ± 0.037 |  | 0.045 ± 0.024 | 0.0267 ± 0.017 |
| **Rickettsiales** | 20.918 ± 23.649 | 36.361 ± 12.894 |  | 3.247 ± 3.221 | 2.321 ± 1.802 |
| **Saccharimonadales** | 0.844 ± 0.577 | 1.269 ± 0.824 |  | 0.141 ± 0.105 | 0.102 ± 0.063 |
| **Solirubrobacterales** | 0.11 ± 0.011 | 0.136 ± 0.092 |  | 0.0227 ± 0.013 | 0.019 ± 0.011 |

**Table S3** % Compositional mean ± standard deviation of dominant bacterial orders in *Daphnia* guts and bodies of *Caullerya-*infected and uninfected samples. Significant differences are highlighted in bold.

| **ZOTU ID** | **Phylum** | **Class** | **Order** | **Family** | **Genus** | **Species** | ***p*sidak** |
| --- | --- | --- | --- | --- | --- | --- | --- |
| ZOTU3 | Proteobacteria | Gammaproteobacteria | Burkholderiales | T34 | Unknown | Unknown | 0.0396 |
| ZOTU1 | Proteobacteria | Gammaproteobacteria | Enterobacteriales | Enterobacteraceae | Unknown | Unknown | 0.0199 |
| ZOTU746 | Proteobacteria | Gammaproteobacteria | Burkholderiales | Chromobacteraceae | *Deefgea* | *Deefgea rivuli* | 0.0199 |
| ZOTU928 | Proteobacteria | Alphaproteobacteria | Reyranellales | Reyranellaceae | *Reyranella* | Unknown | 0.0199 |
| ZOTU407 | Planctomycetota | Planctomycetes | Gemmatales | Gemmataceae | Unknown | Unknown | 0.0485 |
| ZOTU174 | Proteobacteria | Alphaproteobacteria | Rhodobacterales | Rhodobacteriaceae | Unknown | Unknown | 0.0199 |
| ZOTU97 | Planctomycetota | Planctomycetes | Gemmatales | Gemmataceae | Unknown | Unknown | 0.0199 |
| ZOTU927 | Protebacteria | Gammaproteobacteria | Alteromonadales | Shewanellaceae | *Shewanella* | Unknown | 0.0199 |
| ZOTU78 | Planctomycetota | Planctomycetes | Isosphaerales | Isosphaeraceae | Unknown | Unknown | 0.0199 |
| ZOTU367 | Planctomycetota | Phycisphaerae | Phycisphaerales | Phycisphaeraceae | CL500-3 | Unknown | 0.0199 |

**Table S4** ZOTUs identified as Indicator taxa upon comparing *Caullerya-*infected and uninfected *Daphnia* (9999 perm)*. p-*value was adjusted for multiple comparisons using the ‘sidak’ method (“*p*sidak”). See Fig. 4 for abundance of these ZOTUs across samples. Classification presented here follows the NCBI nomenclature.

| **ZOTU** | **Taxonomic Classification** | **Closest known BLAST hit (100% identity)** | **Putative functional ecology** |
| --- | --- | --- | --- |
| ZOTU3 | Order: Burkholderiales; Family: T34 | Uncultured Burkholderiales sp. | The family Burkholderiales is consistently reported as a dominant order in the microbial communities of *Daphnia* sp. (5) and is proposed to have beneficial effects on the host (6). |
| ZOTU1 | Family: Enterobacteraceae; Genus: *Yersiniceae* | *Serratia proteamaculans, Serratia plymuthica, Rahnella aquatilis* and *Yersinia frederiksenii* | *Serratia* sp. are common in aquatic habitats and some have been identified as opportunistic pathogens in humans (7). *S. proteamaculans* has antibiotic properties (8) and can function as a probiotic against the fungal tomato blight (9). |
|  |  |  | *R. aquatilis* is a psychrotolerant pathogen of several aquatic hosts (10, 11). |
|  |  |  | *Y. frederiksenii* is notedly non-pathogenic and can have antibiotic properties against other members of the same genus (12). |
| ZOTU367 | Family: Phycisphaeraceae; Genus: *CL500-3* | Uncultured *Phycisphaera* sp. | *Phycisphaera* sp. form a part of marine biofilms (13) but little is known about their functional ecology (14). |
| ZOTU174 | Phylum: Preteobacteria; Family: Rhodobacteraceae | Uncultured *Rhodobacter* sp. | *Rhodobacter* sp. are autotrophic purple non-sulphur bacteria (15). They can function as probiotics that mitigate lethality caused by the Acute Hepatopancreatic Necrotic Disease (AHPND) in white shrimp (16). |
| ZOTU97 | Phylum: Planctomycetes; Family: Gemmataceae | Uncultured Plactomycetes bacterium | Planctomycetes are unique in their similarity to eukaryotic cellular structure (17). Their presence correlates with cyanobacterial blooms in aquatic ecosystems (18). They form part of aquatic biofilms (13) and are putatively involved in polysaccharide and chitin degradation (19). |
| ZOTU78 | Phylum: Planctomycetes; Family: Isosphaeraceae |  |  |
| ZOTU407 | Phylum: Planctomecetes; Family: Gemmataceae |  |  |
| ZOTU928 | Family: Reyranellaceae; Genus: *Reyranella* | *Reyranella massiliensis* | Biochemical features and enzymatic activity of members of this genus have been characterized, but nothing is yet known about their functional ecology (26, 27). |
| ZOTU927 | Family: Shewanellaceae; Genus: *Shewanella* | Uncultured *Shewanella* sp. | *Shewanella* sp. are physiologically diverse, but typically psychrophilic (28); they are associated with carbon recycling, degradation of organic matter and frequently found in association with aquatic hosts (29). They span the mutualism-parasitism continuum, being reported as pathogenic in eels (30), mollusks (31) and abalones (32), as opportunistic pathogens in human beings (33, 34) as well as probiotics aiding disease resistance in shellfish (35) and flatfish (36). |
| ZOTU746 | Family: Chromobacteriaceae; Species: *Deefgea rivuli* | *Deefgea rivuli* strain IFH1-02IFH1-02 | *Deefgea* strains are found in association with fish (37-40) and mollusks (41) but their functional role is unknown. |

**Table S5** ZOTUs identified as Indicator taxa to differentiate between Caullerya-infected and uninfected Daphnia. The consensus sequence of each ZOTU was BLASTed against the NCBI database, and the closest known bacteria along with their ecology and putative functional capacities are reported here. Taxonomic classification in this table follows NCBI nomenclature, which can differ from Silva database nomenclature. E.g. ZOTU1 (Family: Enterobacteraceae) according to NCBI nomenclature is classified as Family: Yersiniaceae in the Silva database. We chose to use NCBI nomenclature for ZOTUs since this is the most commonly used in literature.
